# Experimental assessment of large mammal population estimates from airborne thermal videography

**DOI:** 10.1101/2024.12.31.630817

**Authors:** Julia S. McElhinny, Gregory D. Larsen, Max Messinger, Chad H. Newbolt, Andrew Whitworth, Stephen S. Ditchkoff, Miles R. Silman, Jared T. Beaver

## Abstract

Wildlife resource management requires reliable, fast, and affordable methods of surveying wildlife populations to develop and adaptively adjust policies. Many methods are used to survey populations of large mammals, and thermal video from drones can yield high rates of detection over large extents with relative speed and safety. In wild populations it can be difficult to estimate detection rates and the accuracy of resulting abundance estimates, especially because many factors of study design and natural systems can influence counts from thermal surveillance. We surveyed a captive white-tailed deer (*Odocoileus virginianus*) population of known size at the 174-ha Auburn Deer Facility in Auburn, AL, USA, and used a designed experiment to investigate the effects of observers, time-of-day, and day-to-day variation on the accuracy of abundance estimates from trial flights using drone-based airborne thermal videography. The experiment consisted of 20 full-census trials, occurring over 4 days, at times-of-day before sunset, after sunset, near midnight, before sunrise, and after sunrise, with each trial video counted by three trained observers. Counts across all trials and observers yielded a mean point estimate of 77 (95% CI 71–83) deer representing 81–97% of the known population range. Flights near sunset yielded estimates of the highest accuracy, within or close to the true range of the population. Variability in estimates was primarily influenced by daily climatic conditions and time-of-day, with only minor observer effects. Results highlight the importance of day-to-day variability in environmental conditions and diel processes, such as nocturnal cooling of the environment and crepuscular peaks in deer activity. The estimates from our experiment and resulting model illustrate that censuses conducted at optimal times-of-day and averaged across multiple days were able to achieve accurate estimates within the known range without correction, and that by averaging across flights, even including suboptimal conditions, thermal videography returned estimates within 4% of the known range with substantial overlap of the estimate. Conversely, single estimates chosen at random from any time-of-day varied widely around the known value, highlighting the need to plan flights with respect to diurnal activity patterns and landscape thermal properties, or, in the absence of information, average multiple surveys across times and days.

## INTRODUCTION

Wildlife management requires accurate and precise estimates of population size, density, and related dynamics to maintain sustainable wildlife populations (Koons et al. 2006). Wildlife biologists use various methods to obtain these estimates, varying greatly in expense, effort, feasibility of deployment, and accuracy, and therefore require evaluation before adoption in management decisions (Collier et al. 2007, Rönnegård et al. 2008). Indirect methods of abundance estimation, such as ungulate pellet surveys, can reduce disturbance by humans during data collection (Camargo-Sanabria and Mandujano 2011, Alves et al. 2013), and other methods, such as mark-and-recapture (Strandgaard 1967), spotlight surveys (Collier et al. 2007), telemetry devices (Duquette et al. 2014, Clement et al. 2015), camera traps (Macauley et al. 2020, Palencia et al. 2021), and more recently environmental DNA (Bohmann et al. 2014), have all been explored as feasible methods to collect population data with varying trade-offs of cost, accuracy, observational scale, and potential disturbance to the target population.

White-tailed deer (*Odocoileus virginianus*; hereafter, deer) are of special management concern because of their high abundance in many regions and related effects on ecosystems, the spread of zoonotic diseases, and economic damages as agricultural and urban pests (Waller and Alverson 1997, Hewitt 2015). Deer are generalists and occupy a wide variety of habitats, including areas occupied by humans, and at high density can degrade forest habitats, altering species composition and reducing tree regeneration by their browsing habits (Waller and Alverson 1997, McShea 2012). Deer and other cervid populations can also transmit diseases to livestock and humans, act as a reservoir for diseases like bovine tuberculosis and brucellosis, and be hosts for zoonotic disease vectors (e.g. ticks, mosquitos) that transmit disease to humans (McShea 2012). Wildlife managers have long regulated hunting harvests to control deer populations (Waller and Alverson 1997, Hewitt 2015), with policies informed by population data, such as estimated abundance, density, and age and sex structure to mitigate the numerous impacts of deer overpopulation (Hansen and Beringer 1997).

Research supporting deer management therefore targets methodologies that provide wildlife managers with population data quickly, reliably and cost-effectively. Aerial surveys are a non-invasive remote sensing method that can rapidly survey deer populations over large areas and reduce some inherent biases that occur in ground-based survey methods (Haroldson et al. 2003, Chabot and Bird 2015, Wang et al. 2019). Extensive research has characterized sources of variability in aerial surveys, including difficulties of locating animals that are nocturnal and animals that are camouflaged in their environment or obscured by a forest canopy (Burgdorf and Weeks 1997, Fleming and Tracey 2008, Shahbazi et al. 2014, Chrétien et al. 2016). Aerial surveys can also yield low accuracy and high variability due to lower sighting probability by human observers compared to ground-based surveys (Pollock and Kendall 1987, Burgdorf and Weeks 1997). Such inaccuracy is attributed to challenges with human visual acuity and observer fatigue when attempting to produce counts from aircraft. Photography and videography can mitigate some of these challenges by recording data for later, more thorough examination (Leedy 1948) or digital image analysis (Chrétien et al. 2016). Nevertheless, many nocturnal and cryptic animals readily evade detection from still photography, videography, or human observers, creating a need for methods for wildlife surveillance beyond the spectra of visible light.

Thermal imaging has been employed in aerial surveys to improve detection beyond that of onboard human observers and to study nocturnally active species (e.g. Havens and Sharp 1998, Witczuk et al. 2018). As large endotherms, deer contrast in the thermal infrared spectrum against many materials and substrates of their environment that might camouflage them in the visible spectrum. This produces a distinct visual signature in thermal imagery, or “thermal signature,” (Croon et al. 1968, Graves et al. 1972) that is characterized by the size, shape and thermal intensity of the animal. The thermal signature facilitates higher detection rates than those from both conventional imagery in the visible light spectrum (Parker Jr. and Driscoll. 1972, Israel 2011, Kissell Jr. and Nimmo. 2011) and from aerial surveys using human observers in real-time (Naugle et al. 1996, Havens and Sharp 1998). However, aerial thermal imaging has been historically uncommon because sensors are often expensive and collect imagery at low resolutions compared to conventional cameras (Naugle et al. 1996), and flight surveys are generally costly and dangerous (Sasse 2003), even more so at night (Watts et al. 2010).

Drones are now a cheaper, safer, and efficient alternative to surveys from occupied aircraft (Chabot and Bird 2015). Drones can collect data at higher spatial and temporal resolutions than many conventional airplane systems because they can be flown safely at low altitudes and in locations, conditions and windows of deployment that are unsuitable or unsafe for larger aircraft (Jones et al. 2006, Hodgson et al. 2018). A variety of platform, sensor and payload options has further revolutionized the collection of environmental data from drones. Because drones can be deployed quickly and with little logistical overhead, they have been deployed over species and habitats where conventional methods are impossible or impractical, increasing monitoring of many diverse species including marine mammals, large terrestrial carnivores, birds, and ungulates (Anderson and Gaston 2013, Chabot and Bird 2015).

Under appropriate conditions, aerial thermal imagery has yielded higher detection rates of deer than spotlight surveys (Naugle et al. 1996, Beaver et al. 2014, Preston et al. 2021) and very high success detecting known and tracked animals (McMahon et al. 2021). But despite these successful applications and a growing number of others, guidelines remain few and uncertain for the collection and interpretation of thermal data. A variety of factors can influence the quality and uncertainty of aerial thermal imagery (**Fig. 1**), some of which may be inherent to the study system—factors associated with the species and its environment—and others of which may be controllable—choice of sensor, flight plan and timing of flight. A confident detection of an animal in thermal imagery depends on a human or algorithmic identification of the animal’s thermal signature amid its environment (Graves et al. 1972), but the contrast between an animal’s signature and its environment is not necessarily consistent across scenarios or even individuals. An animal’s thermal signature is characterized by its visual profile and the amount of thermal energy that its body is releasing into the environment. The visual profile can change with behavior, for example if a deer is bedding down (Wiggers and Beckerman 1993), while thermal brightness can change with physiological shifts, for example when deer seasonally grow thicker fur coats that insulate their thermal energy and reduce contrast with their environment (Croon et al. 1968, Israel 2011). If a study region includes non-focal species with similar thermal signatures, these can be misidentified as the focal species, and daytime surveys with paired color imagery may be necessary to disambiguate species (Larsen et al. 2022), or else thermal video may reveal species-typical movements and behaviors. Structural features of the environment, such as seasonal leaf cover or persistent rock formations, can partially or fully occlude thermal signatures (Naugle et al. 1996, Chrétien et al. 2016). For example, rocks and tree trunks can emit thermal energy and create confounding signatures in video footage (Bernatas and Nelson 2004, Kissell Jr. and Tappe. 2004, McMahon et al. 2021). Survey planning can address some of these concerns by targeting seasons with minimal vegetation, and by surveying between late evening and early morning hours, when thermal contrast is the highest between endotherms and their environments (Bernatas and Nelson 2004, Witczuk et al. 2018). Weather conditions also affect background temperatures and, therefore, the thermal signature of deer; overcast skies can increase detection probability by reflecting and absorbing solar radiation from the top of the atmosphere and thereby preventing the background environment from warming (McMahon et al. 2021). Low temperatures and low fog occurrence also increase the contrast of a thermal signature against its environment (Storm et al. 2011, McMahon et al. 2021), and high winds generally reduce the quality of drone imagery by adding vibration to the aerial platform and camera. Finally, regardless of conditions, parameters of the drone system and flight can affect the quality of data collected. For example, camera resolution and flight altitude affect the ground sample distance, and thereby the resolution of thermal signatures in imagery, and high flight speeds at lower altitudes can cause motion blur and reduce the period of observation over any given animal.

**Figure 1.**
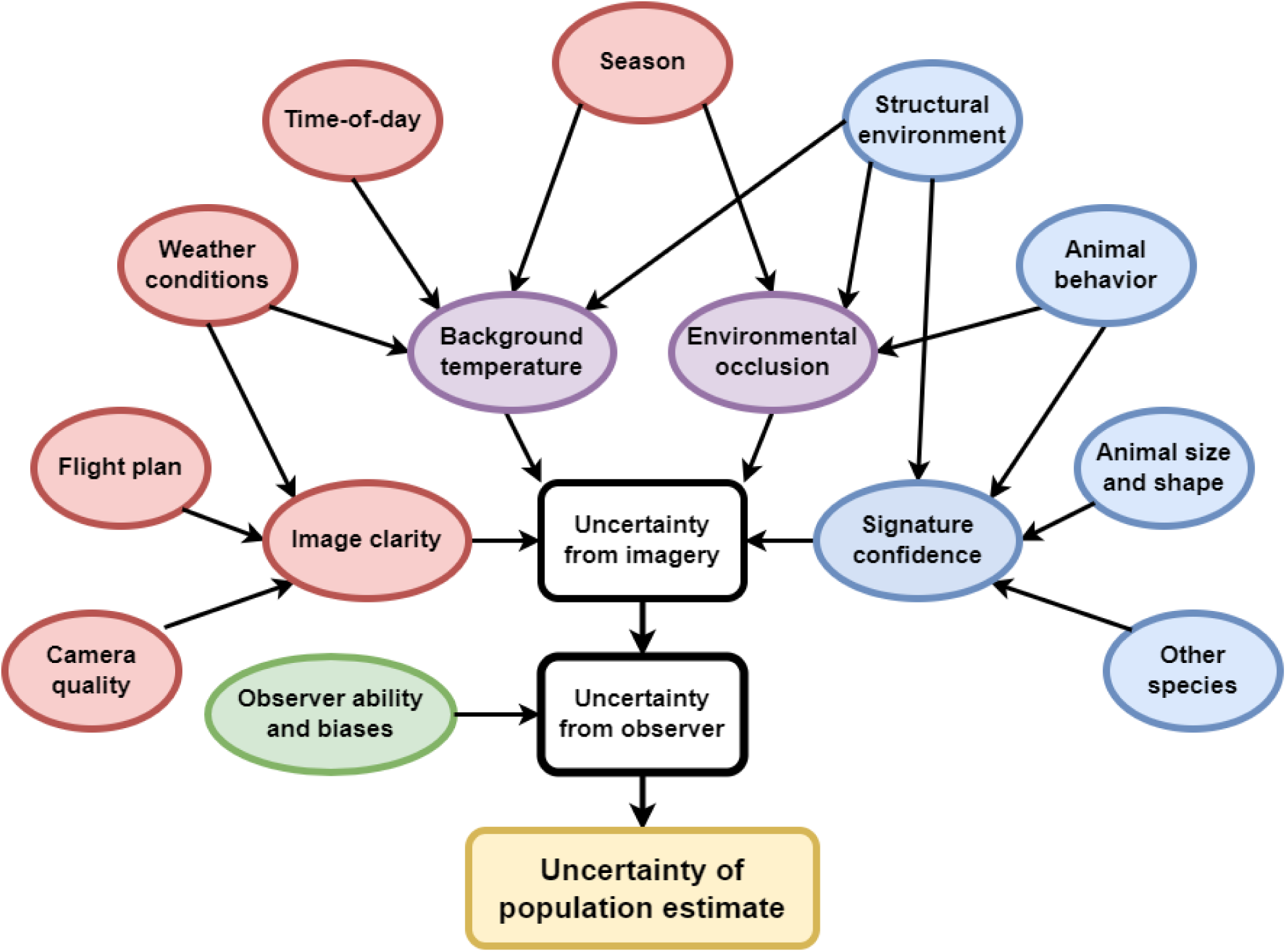
Uncertainty cascades from a variety of sources to affect population estimation from aerial thermal videography with a drone. Sources of uncertainty (ovals) include intrinsic components of the focal population (blue), arising from the species and its ecosystem, and aspects of study design (red) that can often be controlled or selected. Uncertainty from imagery concerns both (1) the detectability of the target species, and (2) the abundance and ambiguity of potential false-positive features. This uncertainty from the imagery is ultimately filtered through the ability and biases of the observer to yield counts from thermal video.

All of these considerations can affect the success of thermal surveys from drones, but they are scarcely studied in combination or to understand their influence on animal detection, species classification, or the accuracy of aggregate abundance estimates.

Abundance estimation is a top concern of wildlife management, and aerial thermal videography must achieve high accuracy as detection rates scale into population estimates if it is to be adopted by practitioners (Haroldson et al. 2003, Storm et al. 2011). Accuracy is a challenging metric, however, because a population’s true size is rarely known (Hodgson et al. 2016). In the absence of a true value, many studies compare estimates from aerial thermal surveys to those from other methods and quantify differences and variation in the method’s detection rate, confounding efforts to standardize methods of accurate population estimation as conditions differ widely between published examples (Burke et al. 2019, Kays et al. 2019). Thermal video has previously achieved density estimates of variable accuracy among mark-recapture surveys, attributed to highly variable detection rates that result from uncertainty in the imagery and uncertainty from the human observers (**Fig. 1**)(Haroldson et al. 2003). Another test of the accuracy of counts of large mammals was focused on seasonal variability and found high accuracy using a small (0.13 km^2^) confined population with short-duration flights using thermal video from a multi-rotor aircraft (Zabel et al. 2023).

Here we assess the accuracy and components of variability in large-scale aerial surveys in heterogeneous landscapes using drone-based thermal videography from a fixed-wing VTOL drone capable of flight times exceeding 1 hr. We surveyed a known, closed population of deer in a designed experiment, with flights spaced every two hours from late afternoon through the night until after sunrise, replicated across four days with varying weather conditions. We analyzed the resulting counts of these trials to estimate the effect of time-of-day, day-to-day variation, and among-observer variation on the ability to recover the known population estimate. These results inform (1) the accuracy of abundance estimation of an ungulate population using aerial thermal videography, (2) the influence of time-of-day, environmental conditions, and observers on the accuracy and agreement of deer detection and counts, and (3) the optimal conditions to maximize reliability of surveys in this system using aerial thermal videography. We also review considerations of study design and provide recommendations for best practices in the use of aerial thermal videography for wildlife population assessments.

## METHODS

### Study area and population

We surveyed a known deer population at Auburn University’s Deer Research Facility in Camp Hill, Alabama (**Fig. 2**) in January 2020. The facility houses 80–95 deer in 1.74 km^2^ of natural habitat enclosed by a 2.5-m tall steel fence. The enclosure is bisected by a large creek and dominated by old pastures, bottomland hardwood trees, and newly planted pines. The deer and other wildlife populations in the enclosure were wild animals captured during construction in October 2007 and their descendants. The population was intensely monitored using annual mark-recapture and camera-trap surveys at the time of this study. The population was not hunted and could move freely throughout the facility (Neuman et al. 2016, Keever et al. 2017, Newbolt et al. 2017, Beaver et al. 2020).

**Figure 2.**
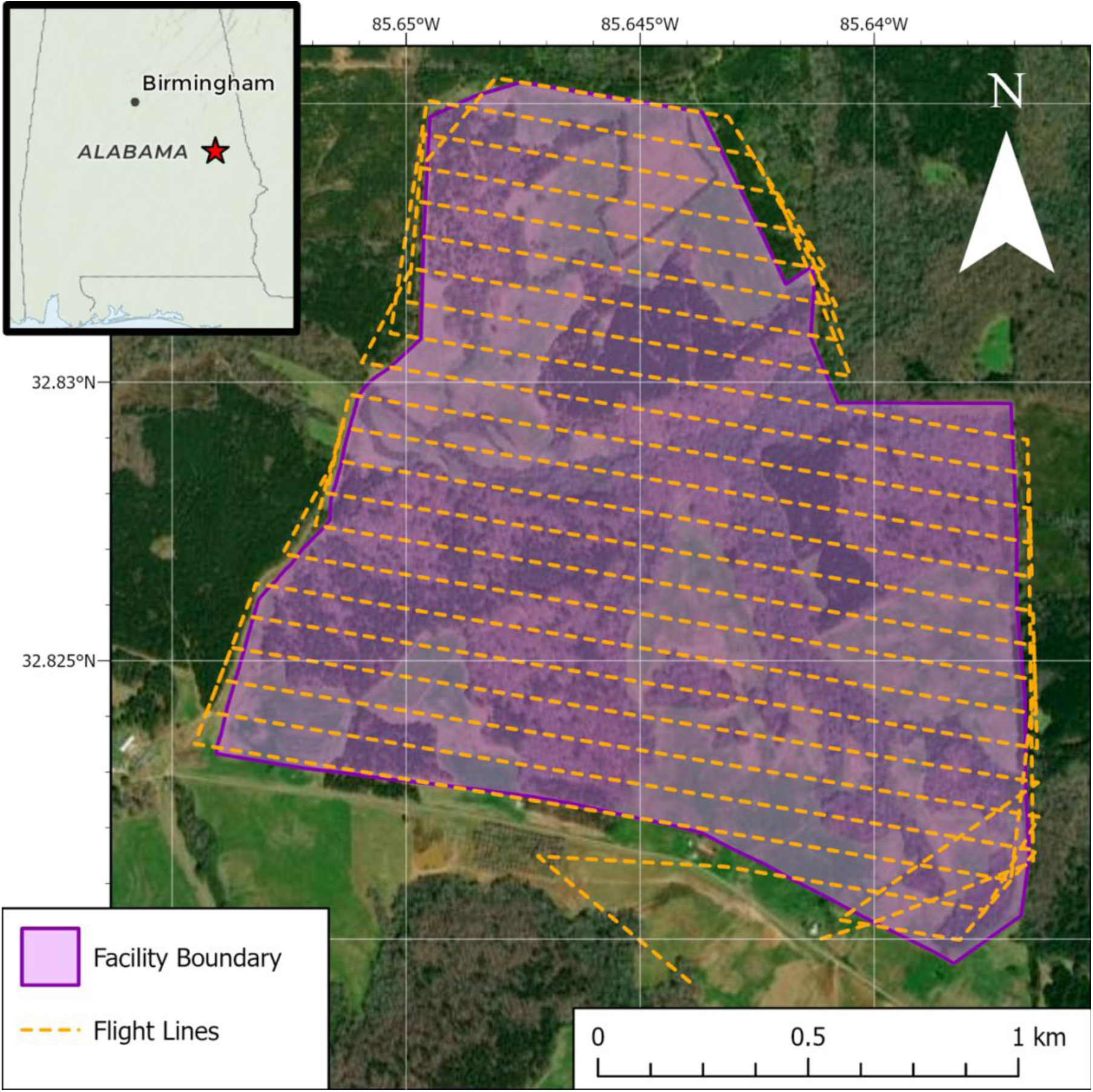
Map of the Auburn Deer Facility in Auburn, AL (red star), as surveyed during January 24–28, 2020. The facility’s enclosed area (purple) was completely surveyed in aerial thermal videography from drones using the same flight plan (orange) over 20 replicate trials to count the deer population under different survey conditions.

### Data Collection

We collected aerial thermal video from a Nimbus VTOL V2 fixed-wing drone (Foxtech, Tianjin, China) using a Cube Autopilot (Hex Technology Ltd, Hong Kong, China) running the ArduPlane software stack from ArduPilot (http://firmware.ardupilot.org/Plane/). We designed flight plans using Mission Planner version 1.3 (http://ardupilot.org/planner/) with terrain-following to ensure a fixed flight altitude. Flights operated at 100 m above ground level (AGL) with a resulting ground sample distance of 13 cm/pixel and horizontal swath of 83.7 m, flying at an average ground speed of 18–19 m/s (ranging 13–26 m/s) and airspeed within 1–2 m/s of the average ground speed. We flew identical replicate flight plans (*n* = 20), each consisting of 24 parallel transects spaced approximately 81.5 m apart, totaling 27.9 km in length, lasting 35 minutes and achieving complete (100%) spatial coverage of the facility (**Fig. 2**).

We collected aerial thermal video using a FLIR Vue Pro R 640 radiometric thermal imager (13-mm lens, 640×512-pixel sensor resolution, 45° horizontal field-of-view, 30 Hz video frame rate, IR spectral band 7.5–13.5 µm; FLIR Systems, Inc., Wilsonville, OR, USA) with simultaneous visible-light (red, green, blue) videography. The thermal imager was self-calibrating and mounted with a nadir view and horizontal camera axis perpendicular to the flight direction. We flew 5 replicate trials each night from the evening of January 24, 2020 to the morning of January 28, 2020, targeting launch times before sunset (∼16:50), after sunset (∼18:20), midnight (∼23:45), before sunrise (∼4:55) and after sunrise (∼6:25), spanning the diel peak of deer activity (Beier and McCollough 1990) and hypothesized to maximize thermal contrast between animal and environment (Chrétien et al. 2016; Witczuk et al. 2018). All launches occurred within 2 minutes of their target times, except the midnight flight which occurred more variably, within 15 minutes of its target time (**Table 1**). Sunset occurred at 17:07– 17:10 on these days, and sunrise occurred at 6:41–6:39. Air temperature and solar irradiation were recorded on a nearby weather gauge (station ID: KALWAVER3, 32.763° N, 85.564° W) approximately 20 km away in Waverly, AL (Weather Underground 2020); air temperatures were recorded every 5 minutes and we averaged these across each trial (**Table 1**). Mean air temperatures (hereafter, temperature) ranged −0.7–10 °C varying by night and time-of-day. Winds were calm (<1 m/s) in all trials except early morning of January 27 which experienced light winds up to 1.5 m/sec. Skies were clear except for night 3 which had cloudy conditions with rain showers occurring near midnight.

**Table 1.** Timing, environmental conditions and weather events of each trial flight. Conditions include temperature (Temp.), humidity (Hum.) and daytime radiance (Rad.). All nights included 5 trials at similar times-of-day: (1) before sunset, (2) after sunset, (3) near midnight, (4) before sunrise, and (5) after sunrise. Count values are averaged across observers (*n* = 3), and bolded values occur within the known range of the deer population (80–95). Sunset occurred at 17:07– 17:10 on these days, and sunrise occurred at 6:41–6:39.

| <u>Night</u> | <u>Date</u> | <u>Launch</u><br><u>time</u> | <u>Temp. (°</u><br><u>C)</u> | <u>Hum.</u><br><u>(%)</u> | <u>Wind</u><br><u>(m/s)</u> | <u>Weather</u> | <u>Count</u> |
| --- | --- | --- | --- | --- | --- | --- | --- |
| 1 | 1-24 | 16:48 | 9.3 | 81.8 | Calm | Clear | **93** |
|  | 1-24 | 18:19 | 6.2 | 90.5 | Calm | Clear | 74 |
|  | 1-24 | 23:35 | 3.8 | 98.8 | Calm | Clear | **81** |
|  | 1-25 | 4:54 | 0.5 | 98.6 | Calm | Clear | 65 |
|  | 1-25 | 6:23 | -0.3 | 99.0 | Calm | Clear | 32 |
| 2 | 1-25 | 16:48 | 7.9 | 74.4 | Calm | Clear | 72 |
|  | 1-25 | 18:18 | 4.9 | 89.3 | Calm | Clear | 71 |
|  | 1-25 | 23:35 | 1.8 | 98.0 | Calm | Clear | 66 |
|  | 1-26 | 4:53 | -0.6 | 99.0 | Calm | Clear | 47 |
|  | 1-26 | 6:23 | -0.7 | 99.0 | Calm | Clear | 64 |
| 3 | 1-26 | 16:54 | 9.9 | 78.3 | Calm | Cloudy | 124 |
|  | 1-26 | 18:17 | 9.0 | 84.0 | Calm | Cloudy | 123 |
|  | 1-26 | 23:47 | 7.5 | 99.0 | Calm | Light Rain | 106 |
|  | 1-27 | 4:53 | 7.0 | 99.0 | NW 1-2 | Cloudy | 126 |
|  | 1-27 | 6:23 | 7.2 | 99.0 | Calm | Cloudy | 65 |
| 4 | 1-27 | 16:48 | 9.4 | 88.5 | Calm | Clear | 69 |
|  | 1-27 | 18:18 | 6.4 | 97.6 | Calm | Clear | **82** |
|  | 1-28 | 00:06 | 1.2 | 99.0 | Calm | Clear | 73 |
|  | 1-28 | 4:54 | -0.2 | 99.0 | Calm | Clear | 68 |
|  | 1-28 | 6:24 | -0.4 | 99.0 | Calm | Clear | 64 |

### Thermal video analysis

Three observers were trained to recognize the characteristic thermal signature of deer in thermal video. The observers then reviewed the video footage from all 20 trials using FLIR tools 6.4 with the default dynamic radiometric scaling to maximize visual contrast in each frame. Observers were blind to flight conditions, known deer population, and counts obtained by other observers. During video review, observers noted occurrences of one or more deer during video recordings, recording a timestamp for each occurrence between the time when deer first appeared in the drone’s field-of-view and the time when deer were no longer present. Observers visually tracked individual deer across frames to ensure that they were not counted more than once. Observers visually identified deer based on the size, shape, and intensity of their thermal signatures relative to background context, static objects and other species (humans, cattle outside the facility boundary), as well as the movement and persistence of these signatures across multiple frames of video. Ambiguous detections were counted or discounted at the discretion of the observer. Each observer’s counts were summed for each trial’s video to yield a population abundance estimate for that trial and observer. We assumed that no deer were counted multiple times per trial, based on the speed of the drone between consecutive transects, the slow locomotive rate of undisturbed deer, and the comparative likelihood of deer entering and exiting transects during the trial. Each observer generated counts of deer from the thermal videography of each trial, yielding 60 counts in total.

### Effects of flight timing and conditions

Data were normally distributed and modeled as count in response to (1) flight times, (2) day, and (3) observer using a linear model with the structure:

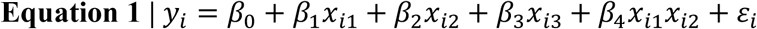

where *μ* describes the count given categorical variables time-of-day (*x*_*i*1_), day (*x*_*i*2_), observer (*x*_*i*3_), and interaction between time-of-day and day (*x*_*i*1_*x*_*i*2_) modified by their respective coefficients (*β*_1–4_) with a fixed intercept (*β*_0_) and variance (*ɛ*_*i*_). We assessed pairwise differences between times-of-day, days, and observers by comparing their estimated marginal means from this model with the Tukey method. Day was modeled as a fixed effect given the low number available for inference (e.g. Gelman and Hill 2007). We conducted all calculations and analyses in R version 4.2.2 (R Core Team 2020), we fit models using the R-package *lme4* (Bates et al. 2015), we analyzed variance of the model and variables using the R-package *car* (Fox and Weisberg 2019, Fox et al. 2023) and we estimated marginal means using the R-package *emmeans* (Searle et al. 1980, Lenth 2023). We report each mean estimate with a 95% confidence interval (95% CI).

## RESULTS

### Thermal videography

Flights yielded variable qualities of video footage for visual assessment, depending on the trial. High-quality thermal videography under optimal conditions (e.g. **Fig. 3a**) described deer with warm thermal signatures against clearly discernible environmental features such as trees, fields, and ponds. Flights under inferior conditions (e.g. **Fig. 3b**) recorded thermal signatures (blue circles) in an attenuated thermal range that created visual ambiguity between deer and other relatively warm features in the environment. Relatively low-quality thermal videography under poor conditions (**Fig. 3c**) yielded thermal signatures (blue circles) with little contrast across the frame to help distinguish deer from other warm features in the environment. Observers noted that high-quality videography was generally collected near sunset, and videography with the poorest quality was collected specifically on night 3 (**Fig. 3c**) amid precipitation and persistently warm temperatures — although deer still contrasted from the background with warmer thermal signatures (**Fig. 3**).

**Figure 3.**
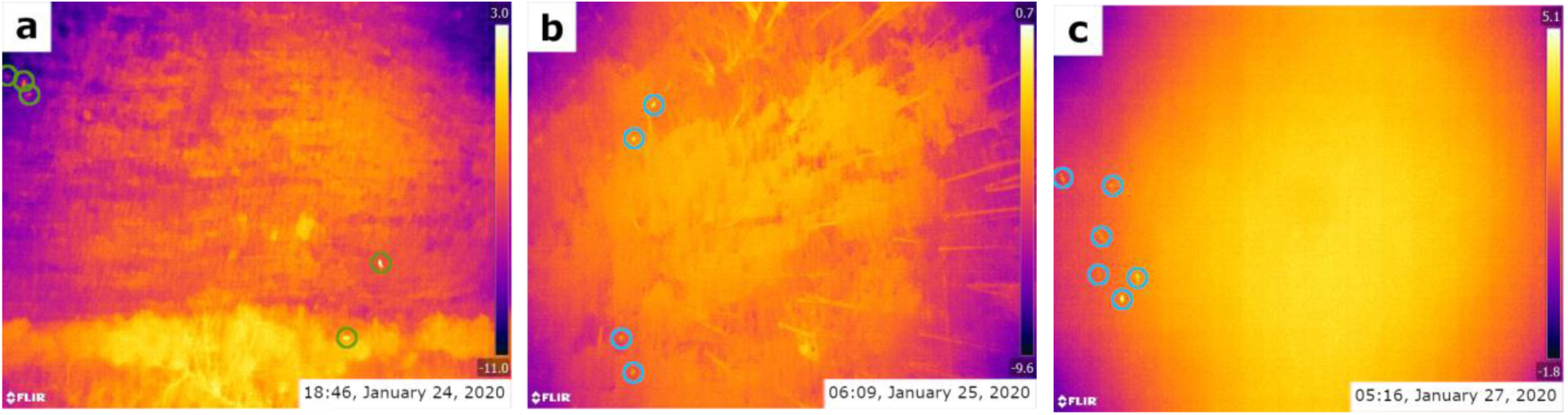
Example frames from thermal videography illustrate visual differences in data quality across nights and times-of-day. High-quality survey conditions (**a**) yield high contrast between deer and background substrates across multiple frames of video, enabling confident identification of deer (green circles) from their background environment. Lower-quality conditions (**b**) create visual ambiguity between deer and environmental features recorded in a similar thermal range (blue circles). Poor conditions (**c**) obscure visual detail among thermal noise, providing little or no context to discriminate between deer and warm environmental features (blue circles). Frames are color-scaled to their maximum and minimum radiometric values (°C, right-side scale). All frames show artifacts of biased radiometry along a center–edge gradient caused by uneven sensor cooling during flight.

### Abundance estimates

Counts from all trials of aerial thermal videography yielded an overall mean abundance estimate of 77.4 (95% CI 71–83) deer, overlapping the known interval of 80–95 and representing 88% (CI 81–95%) of the true abundance interval midpoint of 87.5 and 97% (CI 89–104%) of its lower bound, (**Fig. 4a**). Trials that occurred within 2 hours of sunset most closely estimated the true number of deer with a mean of 89 (95% CI 82–97) across observers and days; trials that occurred within two hours of sunrise yielded an estimate of 66 (95% CI 56–76) when averaged across observers and days (**Figure 4a)**.

**Figure 4.**
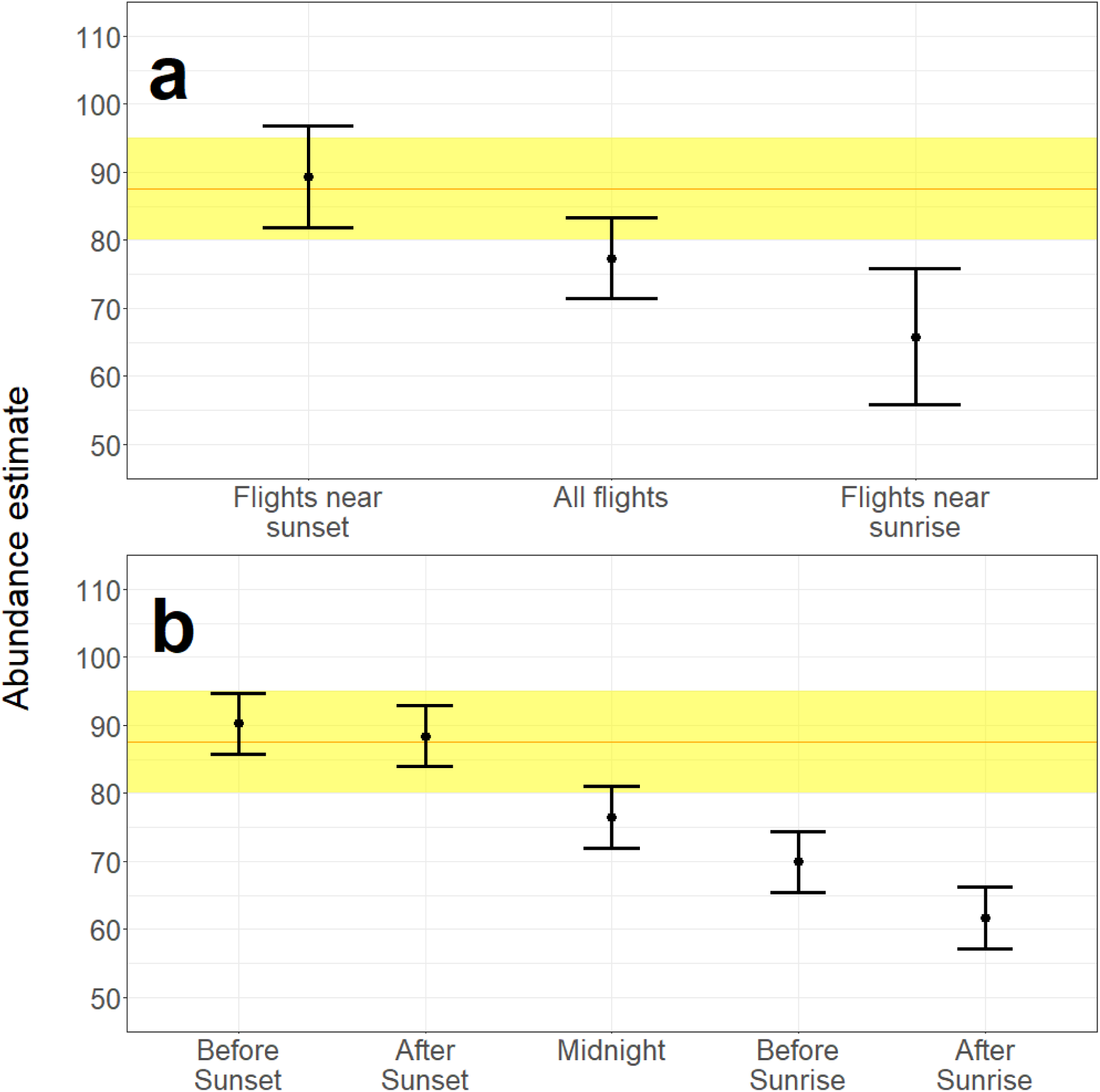
Deer abundance estimates from different times-of-day. Aggregate means (a) describe abundance estimates from flights conducted before or after sunset (left), all flights (center), or flights conducted before or after sunrise (right), averaged across nights and observers. Estimated marginal means (b) describe abundance estimates accounting for fixed effects of observer and day and an interaction of day × time-of-day. Points represent the mean estimates and error bars indicate the 95% CI. Yellow-shaded rectangles indicate the known range of the population (80–95 deer, midpoint 87.5) and orange line indicates the midpoint of that range. Linear modeling yielded population estimates within the known population interval with point estimates within 0.6–3% of the interval midpoint from surveys flown before sunset and after sunset (Fig. 4b).

Abundance estimates were influenced by all modeled effects (**SOM Table 1**). Day explained the most variance (partial *R^2^* = 0.40), followed by the interaction between day × time-of-day (partial *R^2^* = 0.24), time-of-day (partial *R^2^* = 0.23), and with a small effect of observer (*R^2^* = 0.05). Trials that occurred before sunset and after sunset yielded estimates centered in the known range of the population (estimated marginal means of 90 and 88, respectively), while later times-of-day yielded underestimates (**Figure 4b, SOM Figure 1**). Trials that occurred on days 1, 2, and 4 yielded estimates near or below the known range of the population (estimated marginal means of 71, 65, and 71, respectively) but estimates from day 3 overestimated abundance by ∼17% (estimated marginal mean of 102; **SOM Figure 1**). In terms of overall observer performance, one observer yielded an estimated marginal mean within the known range (83), while two yielded estimated marginal means 2.5% and 12.5% below the known range (78, 70), respectively.

## DISCUSSION

Surveys of a known deer population using aerial thermal videography obtained highly accurate estimates with even simple averaging across multiple days and times, giving a point estimate within 11% of the known population midpoint and with a 95% CI overlapping the known range of a closed population. Accounting for when flights occurred, a proxy of both animal behaviors and thermal properties of the landscape, surveys replicated the known interval nearly exactly, with surveys flown near sunset yielding an aggregate mean of 89 deer (95% CI 82–97), compared to a midpoint of 87.5 in a known range of 80–95 deer (**Figs 4 & 5**).

The use of a designed experiment allowed accurate quantification of the components of variation among estimates, with the model capturing 92% of the variability (**SOM Table 1**). Among modeled effects, the day on which surveys took place explained the largest portion of variance in estimates (40%), and we attribute this effect largely to the effects of persistent cloud-cover and precipitation on the effect of radiation exchange between terrestrial objects and the sky (**Table 1**). Atmospheric and surface temperatures, mediated by prior insolation and precipitation, determine the thermal range and contrast within an image, whereas in-air moisture can attenuate atmospheric transmission of thermal light (Bernard et al. 2013, Lloyd 2013). Deer may also show lower activity levels at lower temperatures during winter months (Beier and McCullough 1990), which can decrease their visibility (if bedded down) and detectability (if motionless) in thermal scenes. The effect of thermal changes among days had its largest effect on counts after midnight and through the morning, leading to a large day × time-of-day interaction (24%). The main effect of time-of-day explained a similar proportion of variance (23%) which we attribute to changes in the thermal landscape due to diel cooling and peaks in deer behavior near sunset (Beier and McCullough 1990). Observer effect explained the smallest portion of variance among modeled effects (5%) indicating the technique is robust to differences among observers.

The results from this study suggest employing divergent approaches to dealing with uncertainty, either by averaging across multiple conditions or flights when environmental and subject uncertainty cannot be constrained or by choosing optimal flight times when it can. Both approaches yielded good approximations to true population estimates. We suggest averaging surveys across multiple days to obtain a more accurate estimate across potentially variable weather and animal behavioral conditions. Indeed, our aggregate mean (77) represented a closer approximation at 88% of the known population range midpoint (87.5), than single-day averages across trials of day 1 (71, 81%), day 2 (65, 74%), day 3 (102, 117%), and day 4 (71, 81%). By contrast, we observed that the effects of time-of-day are relatively consistent and predictable across days with counts being accurate near dusk and decreasing relative to the known value at midnight and later surveys (**Fig. 4**), likely due to the environment cooling affecting thermal contrast and decreased deer activity. In this case, we suggest that surveys using aerial thermal videography target times where they capture maximum deer activity, high-contrast thermal imagery, and a low risk of false-positive and false-negative thermal signatures, if possible. Here this corresponds to flights near sunset, but other sites with a high risk of false-positives from target-animal sized objects (as opposed to poor thermal contrast here) might warrant flights near sunrise to prioritize the overnight cooling of inanimate features in the landscape (Garner et al. 1995, Kays et al. 2019, Whitworth et al. 2022).

Aerial thermal imaging entails distinct risks of false-positives and overcounting compared to other methods of surveying deer populations, such as spotlight surveys, camera traps, and aerial surveys using visible-light. Undercounting is a typical bias among abundance estimates of deer from many methods (Forsyth et al. 2022) because deer can camouflage effectively against background substrates or bed down among obscuring vegetation, evading visual detection, especially when they are less mobile. Any survey of mobile animals also includes a risk of counting individuals multiple times as they move in and out of view, but this does not bias estimates if movement is such that the probability of a transect ingress (repeat-count) approximately equals the probability of a transect egress (evaded count) (Buckland et al. 2001). Thermal imagery increases the chance of detecting camouflaged or bedded-down deer, but inanimate objects may also show similar thermal signatures especially if visual context is lacking. We presume that this contributed to the prevailing overestimates on Day 3, possibly in conjunction with higher deer activity (and corresponding detectability) amid the warmer nighttime temperatures. False positives are not generally a special concern in population estimation if they occur consistently across surveys and can be accounted for among other sources of error—indeed, unbiased estimates require them. False positive detections present a rich area for study in thermal imaging, particularly as computer vision and artificial intelligence become part of the aerial imaging work flow. More simply, aerial surveys can be used to locate potentially confounding thermal signatures by identifying their persistent occurrence at recorded locations across similar thermal conditions. Complementary methods, such as collaring and camera traps, can also inform whether variable estimates can be attributed to spatial and temporal changes in deer activity and density.

### Summary and recommendations

This study used a known, closed population of deer to test the efficacy of aerial thermal videography to achieve population censuses. We identified optimal flight times-of-day for the population and demonstrated that thermal videography recovered the true population interval, particularly when sources of variation were modeled, with thermal conditions of the environment and diel patterns of the focal species being especially important. In the absence of such information simple averaging across multiple flights achieved point estimates that were within 12% of the population size range midpoint, and 4% below the lower known range bound. This accuracy emerged even though single-flight estimates reflected a 3-fold variation in population size estimates. These techniques and accuracies should extend to other animals, particularly large mammals, though different focal species and habitats will require different survey planning based on animal behavior, thermal signature, cover type, and the diel trends of ambient temperatures (Burke et al. 2019, Kays et al. 2019, Whitworth et al. 2022). In seasons and environments that yield confounding thermal signatures, such as warm rocks and woody debris of similar size to target species, or thermally attenuated conditions reducing signal-to-noise ratios, false positives are more likely (Graves et al. 1972, Bernatas and Nelson 2004, Storm et al. 2011) and observers might need to identify animals based on movement, if possible, rather than shape and visual contrast alone. These risks are reduced under cloudy daytime conditions, when less solar radiation is available to warm environmental features (McMahon et al. 2021), though this can also deprive observers of helpful context in the thermal scene (Beaver et al. 2020). In all cases, documenting and understanding environmental conditions (temperature, radiation flux, precipitation, aspect) both at the time of flight and from the preceding day can aid interpretation of the resulting estimate. If possible, protocols should investigate and obtain estimated detection rates that are specific to the circumstances of their survey (Kissell Jr. and Tappe. 2004, White 2005, Kissell and Nimmo 2011) and corroborate estimates against those of alternative or conventional methods (Naugle et al. 1996, Baldwin et al. 2023), or at least understand how different components of species biology, physical environment, equipment, and study design contribute to uncertainty and potential bias (**Fig. 1**). Observer bias and interobserver variability had little effect in this study or similar efforts (Beaver et al. 2020), and trained observers were found to be consistent, even across wide variations in average counts (**SOM Fig. 1**; Fleming and Tracey 2008). Future applications of thermal imagery should employ computer vision, as well as incorporate radiometric analysis to improve accuracy, improve replicability, and accelerate video analysis (Christiansen et al. 2014, Ward et al. 2016, Seymour et al. 2017). Thermal surveillance is an accurate and robust means of estimating and characterizing deer and other wildlife populations, poised to improve further with continued development high-definition sensors, fusion between thermal and available visual data, incorporating movement, incorporating locations of animals in previous imagery to prevent false positives, and the application of artificial intelligence (He et al. 2021, Whitworth et al. 2022, Bao et al. 2023).

## Notes

### Competing Interest Statement

The authors have declared no competing interest.

## REFERENCES

Alves, J., A. A. Silva, A. M. V. M. Soares, and C. Fonseca. 2013. Pellet group count methods to estimate red deer densities: precision, potential accuracy and efficiency. Mammalian Biology 78:134–141.

Anderson, K., and K. J. Gaston. 2013. Lightweight unmanned aerial vehicles will revolutionize spatial ecology. Frontiers in Ecology and the Environment 11:138–146.

Baldwin, R. W., J. T. Beaver, M. Messinger, J. Muday, M. Windsor, G. D. Larsen, M. R. Silman, and T. M. Anderson. 2023. Camera Trap Methods and Drone Thermal Surveillance Provide Reliable, Comparable Density Estimates of Large, Free-Ranging Ungulates. Animals 13:1884.

Bao, F., X. Wang, S. H. Sureshbabu, G. Sreekumar, L. Yang, V. Aggarwal, V. N. Boddeti, and Z. Jacob. 2023. Heat-assisted detection and ranging. Nature 619:743–748.

Bates, D., M. Mächler, B. Bolker, and S. Walker. 2015. Fitting Linear Mixed-Effects Models Using lme4. Journal of Statistical Software 67:1–48.

Beaver, J. T., R. W. Baldwin, M. Messinger, C. H. Newbolt, S. S. Ditchkoff, and M. R. Silman. 2020. Evaluating the Use of Drones Equipped with Thermal Sensors as an Effective Method for Estimating Wildlife. Wildlife Society Bulletin 44:434–443.

Beaver, J. T., C. A. Harper, R. E. Kissell Jr., L. I. Muller, P. S. Basinger, M. J. Goode, F. T. Manen, W. Winton, and M. L. Kennedy. 2014. Aerial vertical-looking infrared imagery to evaluate bias of distance sampling techniques for white-tailed deer. Wildlife Society Bulletin 38:419–427.

Beier, P., and D. R. McCullough. 1990. Factors Influencing White-Tailed Deer Activity Patterns and Habitat Use. Wildlife Monographs 3–51.

Bernard, E., N. Riviere, M. Renaudat, P. Guiset, M. Pealat, and E. Zenou. 2013. Experiments and models of active and thermal imaging under bad weather conditions. Pages 14–25 in. Electro-Optical Remote Sensing, Photonic Technologies, and Applications VII; and Military Applications in Hyperspectral Imaging and High Spatial Resolution Sensing. Volume 8897. SPIE.

Bernatas, S., and L. Nelson. 2004. Sightability model for California bighorn sheep in canyonlands using forward-looking infrared (FLIR. Wildlife Society Bulletin 32:638–647.

Bohmann, K., A. Evans, M. T. P. Gilbert, G. R. Carvalho, S. Creer, M. Knapp, D. W. Yu, and M. Bruyn. 2014. Environmental DNA for wildlife biology and biodiversity monitoring. Trends in Ecology and Evolution 29:358–367.

Buckland, S. T., D. R. Anderson, K. P. Burnham, J. L. Laake, D. L. Borchers, and L. Thomas. 2001. Introduction to Distance Sampling: Estimating Abundance of Biological Populations. Oxford University Press.

Burgdorf, S. J., and H. P. Weeks Jr. 1997. Aerial censusing of white-tailed deer and comparison to sex-age-kill population estimates in Northern Indiana. Proceedings of the Indiana Academy of Science 106:201–211.

Burke, C., M. Rashman, S. Wich, A. Symons, C. Theron, and S. Longmore. 2019. Optimizing observing strategies for monitoring animals using drone-mounted thermal infrared cameras. International Journal of Remote Sensing 40:439–467.

Camargo-Sanabria, A. A., and S. Mandujano. 2011. Comparison of pellet-group counting methods to estimate population density of white-tailed deer in a Mexican tropical dry forest. Tropical Conservation Science 4:230–243.

Chabot, D., and D. M. Bird. 2015. Wildlife research and management methods in the 21st century: Where do unmanned aircraft fit in? Journal of Unmanned Vehicle Systems 3:137–155.

Chrétien, L.-P., J. Théau, and P. Ménard. 2016. Visible and thermal infrared remote sensing for the detection of white-tailed deer using an unmanned aerial system. Wildlife Society Bulletin 40:181–191.

Christiansen, P., K. A. Steen, R. N. Jørgensen, and H. Karstoft. 2014. Automated Detection and Recognition of Wildlife Using Thermal Cameras. Sensors 14:13778–13793.

Clement, M. J., J. M. O’Keefe, and B. Walters. 2015. A method for estimating abundance of mobile populations using telemetry and counts of unmarked animals. Ecosphere 6:1–13.

Collier, B. A., S. S. Ditchkoff, J. B. Raglin, and J. M. Smith. 2007. Detection Probability and Sources of Variation in White-Tailed Deer Spotlight Surveys. The Journal of Wildlife Management 71:277–281.

Croon, G. W., D. R. McCullough, C. E. Olson Jr., and L. M. Queal. 1968. Infrared scanning techniques for big game censusing. Journal of Wildlife Management 32:751–759.

Duquette, J. F., J. L. Belant, N. J. Svoboda, D. E. Beyer Jr., and C. A. Albright. 2014. Comparison of occupancy modeling and radiotelemetry to estimate ungulate population dynamics. Population Ecology 56:481–492.

Fleming, P. J. S., and J. P. Tracey. 2008. Some human, aircraft and animal factors affecting aerial surveys: how to enumerate animals from the air. Wildlife Research 35:258–267.

Forsyth, D. M., S. Comte, N. E. Davis, A. J. Bengsen, S. D. Côté, D. G. Hewitt, N. Morellet, and A. Mysterud. 2022. Methodology matters when estimating deer abundance: a global systematic review and recommendations for improvements. The Journal of Wildlife Management 86:e22207.

Fox, J., and S. Weisberg. 2019. An R Companion to Applied Regression. Third. Sage, Thousand Oaks CA.

Fox, J., S. Weisberg, B. Price, D. Adler, D. Bates, G. Baud-Bovy, B. Bolker, S. Ellison, D. Firth, M. Friendly, G. Gorjanc, S. Graves, R. Heiberger, P. Krivitsky, R. Laboissiere, M. Maechler, G. Monette, D. Murdoch, H. Nilsson, D. Ogle, B. Ripley, T. Short, W. Venables, S. Walker, D. Winsemius, A. Zeileis, and R-Core. 2023. car: Companion to Applied Regression. <https://cran.r-project.org/web/packages/car/index.html>. Accessed 8 May 2024.

Garner, D. L., H. B. Underwood, and W. F. Porter. 1995. Use of modern infrared thermography for wildlife population surveys. Environmental Management 19:233–238.

Graves, H. B., E. D. Bellis, and W. M. Knuth. 1972. Censusing white-tailed deer by airborne thermal infrared imagery. Journal of Wildlife Management 36:875–884.

Hansen, L., and J. Beringer. 1997. Managed hunts to control white-tailed deer populations on urban public areas in Missouri. Wildlife Society Bulletin 25:484–487.

Haroldson, B. S., E. P. Wiggers, J. Beringer, L. P. Hansen, and J. B. McAnich. 2003. Evaluation of aerial thermal imaging for detecting white-tailed deer in a deciduous forest environment. Wildlife Society Bulletin 31:1188–1197.

Havens, K. J., and E. J. Sharp. 1998. Using thermal imagery in the aerial survey of animals. Wildlife Society Bulletin 26:17–23.

He, Y., B. Deng, H. Wang, L. Cheng, K. Zhou, S. Cai, and F. Ciampa. 2021. Infrared machine vision and infrared thermography with deep learning: A review. Infrared Physics & Technology 116:103754.

Hewitt, D. G. 2015. Hunters and the conservation and management of white-tailed deer (Odocoileus virginianus. International Journal of Environmental Studies 72:839–849.

Hodgson, J. C., S. M. Baylis, R. Mott, A. Herrod, and R. H. Clarke. 2016. Precision wildlife monitoring using unmanned aerial vehicles. Scientific Reports 6:22574.

Hodgson, J. C., R. Mott, S. M. Baylis, T. T. Pham, S. Wotherspoon, A. D. Kilpatrick, R. Raja Segaran, I. Reid, A. Terauds, and L. P. Koh. 2018. Drones count wildlife more accurately and precisely than humans. Methods in Ecology and Evolution 9:1160–1167.

Israel, M. 2011. A UAV-based roe deer fawn detection system. International Archives of the Photogrammetry, Remote Sensing, and Spatial Information Sciences 38:51–55.

Jones, G. P., IV, L. G. Pearlstine, and H. F. Percival. 2006. An assessment of small unmanned aerial vehicles for wildlife research. Wildlife Society Bulletin 34:750–758.

Kays, R., J. Sheppard, K. Mclean, C. Welch, C. Paunescu, V. Wang, G. Kravit, and M. Crofoot. 2019. Hot monkey, cold reality: surveying rainforest canopy mammals using drone-mounted thermal infrared sensors. International Journal of Remote Sensing 40:407–419.

Keever, A. C., C. P. McGowan, S. S. Ditchkoff, P. K. Acker, J. B. Grand, and C. H. Newbolt. 2017. Efficacy of N-mixture models for surveying and monitoring white-tailed deer populations. Mammal Research 62:413–422.

Kissell Jr., R. E., and S. K. Nimmo. 2011. A technique to estimate white-tailed deer Odocoileus virginianus density using vertical-looking infrared imagery. Wildlife Biology 17:85–92.

Kissell Jr., R. E., and P. A. Tappe. 2004. Assessment of thermal infrared detection rates using white-tailed deer surrogates. Journal of the Arkansas Academy of Science 58:70–73.

Kissell, R. E., and S. K. Nimmo. 2011. A technique to estimate white-tailed deer Odocoileus virginianus density using vertical-looking infrared imagery. Wildlife Biology 17:85–92.

Koons, D. N., R. F. Rockwell, and J. B. Grand. 2006. Population Momentum: Implications for Wildlife Management. The Journal of Wildlife Management 70:19–26.

Larsen, G. D., A. C. Seymour, E. L. Richmond, L. M. Divine, E. E. Moreland, E. Newton, J. M. London, and D. W. Johnston. 2022. Drones reveal spatial patterning of sympatric Alaskan pinniped species and drivers of their local distributions. Drone Systems and Applications 10:235–255.

Leedy, D. L. 1948. Aerial Photographs, Their Interpretation and Suggested Uses in Wildlife Management. The Journal of Wildlife Management 12:191–210.

Lenth, R. V. 2023. emmeans: Estimated Marginal Means, aka Least-Squares Means. <https://CRAN.R-project.org/package=emmeans>.

Lloyd, J. M. 2013. Thermal Imaging Systems. Springer Science & Business Media.

Macauley, L. T., R. Sollmann, and R. H. Barrett. 2020. Estimating deer populations using camera traps and natural marks. Journal of Wildlife Management 84:301–310.

McMahon, M. C., M. A. Ditmer, E. J. Isaac, S. A. Moore, and J. D. Forester. 2021. Evaluating unmanned aerial systems for the detection and monitoring of moose in northeastern Minnesota. Wildlife Society Bulletin 45:312–324.

McShea, W. J. 2012. Ecology and management of white-tailed deer in a changing world. Annals of the New York Academy of Sciences 1249:45–56.

Naugle, D. E., J. A. Jenks, and B. J. Kernohan. 1996. Use of thermal infrared sensing to estimate density of white-tailed deer. Wildlife Society Bulletin 24:37–43.

Neuman, T. J., C. H. Newbolt, S. S. Ditchkoff, and T. D. Steury. 2016. Microsatellites reveal plasticity in reproductive success of white-tailed deer. Journal of Mammalogy 97:1441–1450.

Newbolt, C. H., S. Rankin, and S. S. Ditchkoff. 2017. Temporal and Sex-related Differences in use of Baited Sites by White-tailed Deer. Journal of the Southeastern Association of Fish and Wildlife Agencies 4:109–114.

Palencia, P., J. M. Rowcliffe, J. Vincente, and P. Acevedo. 2021. Assessing camera trap methodologies used to estimate density of unmarked populations. Journal of Applied Ecology 58:1583–1592.

Parker Jr., H. D., and R. S. Driscoll. 1972. An experiment in deer detection by thermal scanning. Journal of Range Management 25:480–481.

Pollock, K. H., and W. L. Kendall. 1987. Visibility bias in aerial surveys: a review of estimation procedures. Journal of Wildlife Management 51:502–510.

Preston, T. M., M. L. Wildhaber, N. S. Green, J. L. Albers, and G. P. Debenedetto. 2021. Enumerating white-tailed deer using unmanned vehicles. Wildlife Society Bulletin 45:97–108.

R Core Team. 2020. R: A Language and Environment for Statistical Computing. R Foundation for Statistical Computing, Vienna, Austria.

Rönnegård, L., H. Sand, H. Andrén, J. Månsson, and A. Pehrson. 2008. Evaluation of four methods used to estimate population density of moose Alces alces. Wildlife Biology 14:358–371.

Sasse, D. B. 2003. Job-Related Mortality of Wildlife Workers in the United States, 1937-2000. Wildlife Society Bulletin (1973-2006) 31:1015–1020.

Searle, S. R., F. M. Speed, and G. A. Milliken. 1980. Population Marginal Means in the Linear Model: An Alternative to Least Squares Means. The American Statistician 34:216–221.

Seymour, A. C., J. Dale, M. Hammill, P. N. Halpin, and D. W. Johnston. 2017. Automated detection and enumeration of marine wildlife using unmanned aircraft systems (UAS) and thermal imagery. Scientific Reports 7:45127.

Shahbazi, M., J. Théau, and P. Ménard. 2014. Recent applications of unmanned aerial imagery in natural resource management. GIScience and Remote Sensing 51:339–365.

Storm, D. J., M. D. Samuel, T. R. Deelen, K. D. Malcolm, R. E. Rolley, N. A. Frost, D. P. Bates, and B. J. Richards. 2011. Comparison of visual-based helicopter and fixed-wing forward-looking infrared surveys for counting white-tailed deer Odocoileus virginianus. Wildlife Biology 17:431–440.

Strandgaard, H. 1967. Reliability of the Peterson method tested on a roe-deer population. Journal of Wildlife Management 31:643–651.

Waller, D. M., and W. S. Alverson. 1997. The White-Tailed Deer: A Keystone Herbivore. Wildlife Society Bulletin (1973-2006) 25:217–226.

Wang, D., Q. Shao, and H. Yue. 2019. Surveying wild animals from satellites, manned aircraft and unmanned aerial systems (UASs): a review. Remote Sensing 11.

Ward, S., J. Hensler, B. Alsalam, and L. F. Gonzalez. 2016. Autonomous UAVs wildlife detection using thermal imaging, predictive navigation and computer vision. Pages 1–8 in. 2016 IEEE Aerospace Conference.

Watts, A. C., J. H. Perry, S. E. Smith, M. A. Burgess, B. E. Wilkinson, Z. Szantoi, P. G. Ifju, and H. F. Percival. 2010. Small unmanned aircraft systems for low-altitude aerial surveys. Journal of Wildlife Management 74:1614–1619.

Weather Underground. 2020. Weather History for KALWAVER3. <https://www.wunderground.com/dashboard/pws/KALWAVER3>.

White, G. C. 2005. Correcting wildlife counts using detection probabilities. Wildlife Research 32:211–216.

Whitworth, A., C. Pinto, J. Ortiz, E. Flatt, and M. Silman. 2022. Flight speed and time of day heavily influence rainforest canopy wildlife counts from drone-mounted thermal camera surveys. Biodiversity and Conservation 31:3179–3195.

Wiggers, E. P., and S. F. Beckerman. 1993. Use of thermal infrared sensing to survey white-tailed deer populations. Wildlife Society Bulletin 21:263–268.

Witczuk, J., S. Pagacz, A. Zmarz, and M. Cypel. 2018. Exploring the feasibility of unmanned aerial vehicles and thermal imaging for ungulate surveys in forests – preliminary results. International Journal of Remote Sensing 39:5504–5521.

Zabel, F., M. A. Findlay, and P. J. C. White. 2023. Assessment of the accuracy of counting large ungulate species (red deer Cervus elaphus) with UAV-mounted thermal infrared cameras during night flights. Wildlife Biology 2023:e01071.

