## Supplemental Online Material for "Experimental assessment of large mammal population estimates from airborne thermal videography"

**SUPPORTING ONLINE MATERIAL:** McElhinny et al.

**SOM Table 1 |** Analysis of variance table for linear model estimating abundance as a function of fixed effects, day, time-of-day, and observer, and an interaction of day × time-of-day. Day explained the most variability in estimates, followed by an interaction of day × time-of-day, time-of-day, and finally observer. The model had an overall ratio of explained variance ($R^2$) of 0.92.

| | Type-III sum of squares | *Partial* $R^2$ | df | F-value | *P*-value |
|---|---|---|---|---|---|
| Day | 12527 | 0.40 | 3 | 70.4 | $1.4\times 10^{-15}$ |
| Time-of-day | 7090 | 0.23 | 4 | 29.9 | $2.9\times 10^{-11}$ |
| Observer | 1587 | 0.05 | 2 | 13.4 | $4.0\times 10^{-05}$ |
| Day × Time-of-day | 7535 | 0.24 | 12 | 10.6 | $9.5\times 10^{-09}$ |

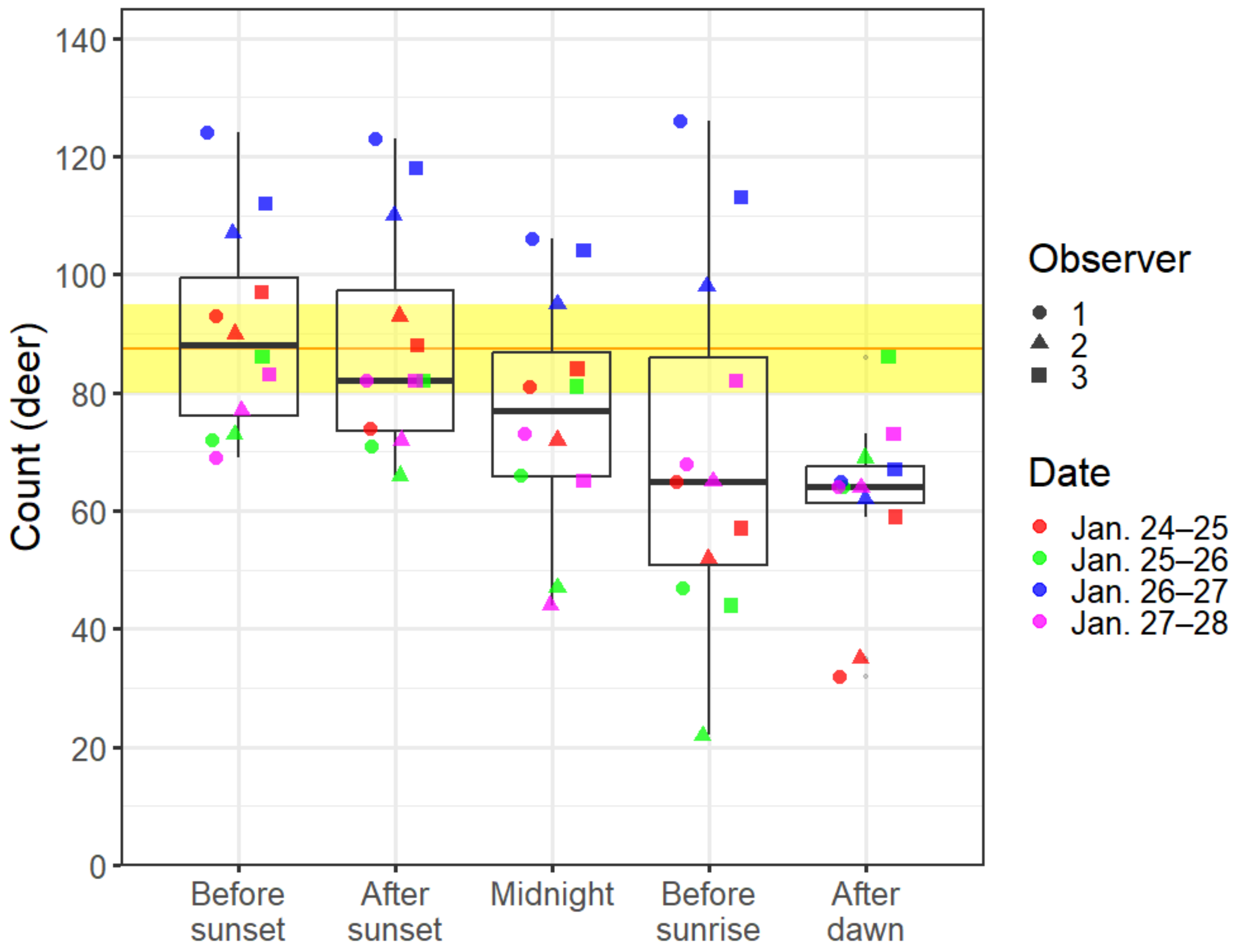


**SOM Figure 1** | Counts of deer grouped by time-of-day. Boxplots represent the distribution of estimates across nights and observers for each flight window. Individual counts are overlaid and distinguished by night (colors) and observer (shapes). Each box represents the IQR; horizontal lines within the boxes represent median values; whiskers represent maximum values (+1.5 IQR) and minimum values (-1.5 IQR); and outliers are plotted beyond the whiskers. The yellow-shaded rectangle indicates the known range of the population (80–95 deer) and the orange line represents the midpoint of that range.
